# Identification and mapping of human lymph node stromal cell subsets by combining single-cell RNA sequencing with spatial transcriptomics

**DOI:** 10.1101/2023.08.18.553530

**Authors:** Cristoforo Grasso, Janna Roet, Catarina Gago de Graça, Johanna F. Semmelink, Ester Remmerswaal, Aldo Jongejan, Perry D. Moerland, Reina E. Mebius, Lisa G. M. van Baarsen

## Abstract

Lymph node stromal cells (LNSCs) have a crucial immunomodulatory function, but their heterogeneity in human is incompletely understood. Here, we report the single cell RNA sequencing (scRNA-seq) of 12’000 LNSCs isolated from a human lymph node (LN). This study comprehensively defines the gene signatures of 10 fibroblast subtypes: CCL21^+^SC, CCL19^+^SC, CD34^+^CXCL14^+^SC, pericytes, DES^+^SC, LAMP5^+^SC, NR4A1^+^BCAM^+^ SC, HLA-DR^+^SC, SEPT4^+^SC and GLDN^+^SC. To explore the heterogeneous stromal compartment within the complex LN tissue architecture, we integrated the scRNA-seq profiles of the identified LNSC subsets with a publicly available human spatial transcriptomic LN dataset and predicted their location within the complex LN tissue architecture. Each LNSC subtype was spatially restricted to specific LN regions, indicating different LNSC-lymphocyte interactions which was further investigated using NicheNet. The positioning of distinct LNSC subtypes in different LN regions sets the stage for future research on the relationship between LNSC-specific niches and immunomodulatory function during health and disease.

## Introduction

Adaptive immune responses are initiated in lymphoid organs, where immune cells surveil to protect our body against pathogens or tumour progression^1^. In a lymph node, non-hematopoietic (CD45-) and hematopoietic cells (CD45+) coexist to orchestrate immunity. The non-hematopoietic cells make up approximately 5% of the total cells of a lymph node^2^. These non-hematopoietic cells, also known as lymph node stromal cells (LNSCs) provide structural and mechanic support to the lymph node but also modulate immune cell maturation, migration and activation^3^. Two major populations of LNSCs can be distinguished, namely endothelial (CD31+) cells, further subdivided into blood (BECs) and lymphatic endothelial cells (LECs), and non-endothelial (CD31-) cells, which are subdivided into fibroblast reticular cells (FRCs) and double negative (DN) cells^4–8^. Lymphocytes can access the lymph node from the blood via high endothelial cells, which form venules and support the multistep leukocyte extravasation cascade^9^. Lymphatic vessels carry self- and non-self-antigens and multiple types of immune cells to the draining lymph node, for initiating immune activation or induction of peripheral tolerance^10^. FRCs secrete components of the reticular network and regulate traffic of lymphocytes and dendritic cells (DCs) over the network^11^. Additionally, LNSCs can present antigens,^12,13^ attract lymphocytes,^14,15^ and B-cells^16^ thereby mediating peripheral immune tolerance^17,18^.

Technological advances, such as single-cell RNA sequencing (scRNA-seq) have expanded our understanding of LNSC heterogeneity^5,19–24^, and how heterogeneity may impact LNSCs immune modulation functions^25^. Heterogeneity of LNSCs reflects the variety of interactions and synergy with the immune system within secondary lymphoid organs. Nowadays, it is clear that dysfunction or depletion of human LNSCs results in a severely impaired immune response^26,27,28^. Therefore, it is of paramount importance to investigate the role of human stromal cells in their LN microenvironment as this may lead to the discovery of potential targets for immunomodulation. The current study employs scRNA-seq and spatial transcriptomics to assess the heterogeneity of human LNSCs, and to identify in which areas of the lymph node each subset is located. With our approach we found ten clusters of fibroblastic cells: NR4A1+BCAM+ SC, CCL21+ SC, CCL19+ SC, CD34+ CXCL14+SC, pericytes, DES+ SC, LAMP5+ SC, HLA-DR+ SC, SEPT4+ SC, and GLDN+ SC. Next, we predicted the locations of each subset in the lymph node by integrating our single-cell RNA profiles with publicly available spatial transcriptomics data. Immunofluorescence staining on lymph node sections was used to validate the predicted location of the newly discovered GLDN+ SC in lymph node follicles. Furthermore, NicheNet was applied to reveal potential T- and B-cell interactions with different LNSC subtypes.

## Results

### Single-cell RNA sequencing analysis of human lymph node stromal cells

To reveal the heterogeneity and molecular blueprint of human LNSCs we have enzymatically digested a human lymph node to obtain a single cell suspension. Subsequently, 4 major human peripheral lymph node CD45− cell subtypes were sorted and processed for droplet-based scRNAseq (10x Chromium) (Fig. 1a, a detailed report of our sorting strategy is included in the Supplementary Note Fig.S1a). Hence, endothelial and non-endothelial LNSCs were classified as CD45-CD235a- and further divided into the four major LNSC subtypes based on the expression of PDPN and CD31: FRC (PDPN+, CD31-), DN (PDPN-, CD31-), LECs (PDPN+, CD31+), and BECs (PDPN-, CD31+). Accordingly, viable lymph node stromal cells were enriched by fluorescence-activated cell sorting (FACS) and we ensured that our isolated stromal cell suspension contained cells from all major LNSC subtypes (Fig. 1b). Sorted cells were sequenced with high coverage using 10x Chromium protocol. After data pre-processing, we performed principal component analysis (PCA) followed by unsupervised clustering visualized on Uniform Manifold Approximation and Projection (UMAP) for visualization of 9,267 clustered cells using a graph-based method. Based on UMAP reduction three distinct cell types (fibroblasts, BECs, LECs) could be clearly separated (Fig. 1c). Based on expression of canonical markers^17,25,26^ we could annotate 7,122 fibroblasts expressing *PDGFRA* and/or *PDGFRB* representing the largest group of our dataset (Figure 1d), 1’244 BECs positive for *PECAM1* and *VWF* and 901 LECs expressing *LYVE1* and/or *PROX1* (Fig. 1e).

**Figure 1.**
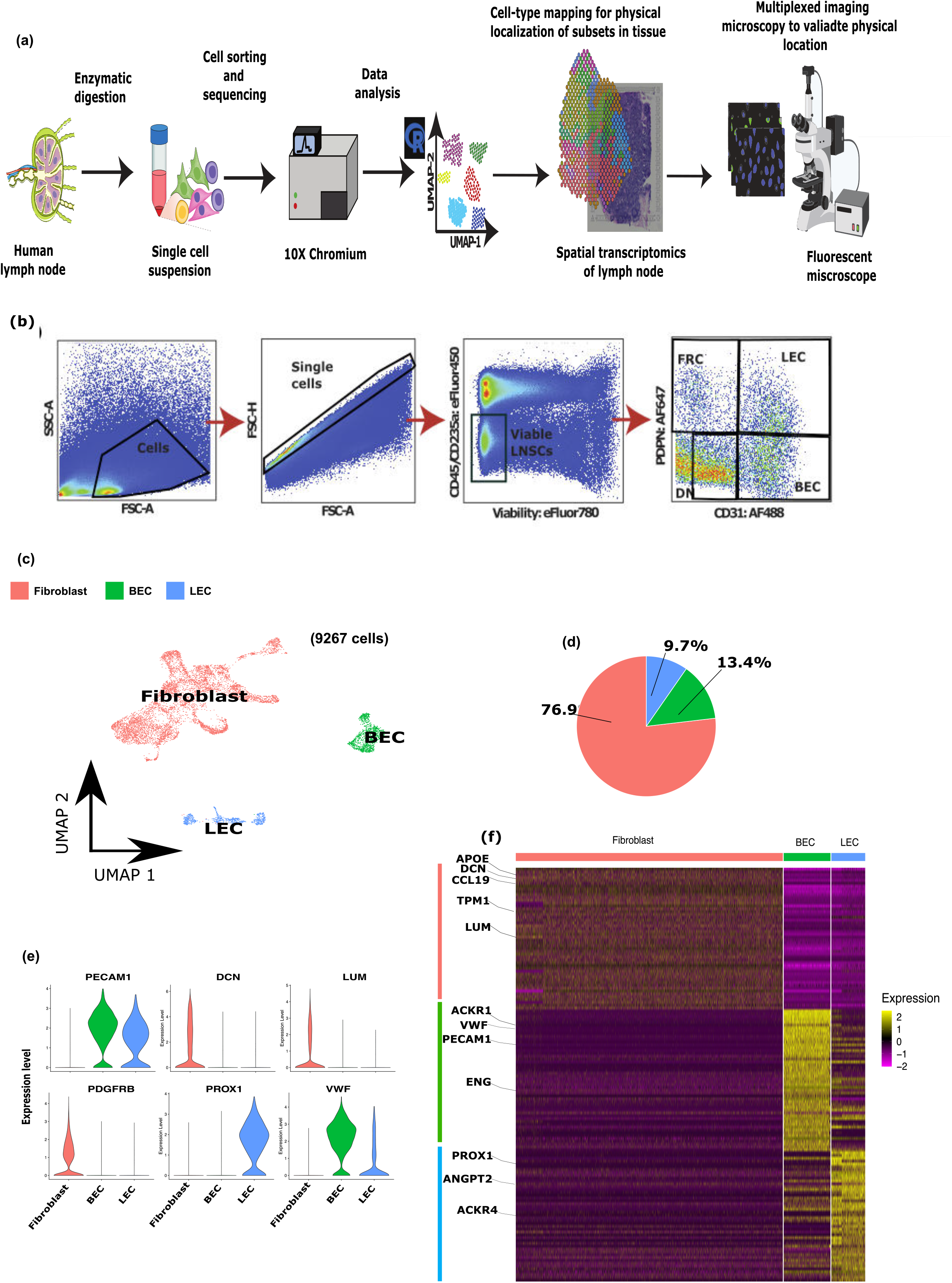
Experimental design and scRNA-Seq identify the major LNSC populations. **a.** Schematic representation of experimental design and techniques used in this study. **b.** Flow cytometry gating strategy used to sort viable CD45- CD235a- single cell containing DN, FRC, LEC and BEC (see Figure S1 for more details). **c.** Two- dimensional clustering of 9,267 LNSCs analysed by scRNAseq. LNSCs are color- coded based on their main phenotype: fibroblasts (red), LEC (blue), BEC (green). **d.** Pie chart showing frequency on each phenotype within the total LNSCs analysed. **e.** Violin plots showing log-normalized expression levels of marker genes for each phenotype: *PECAM1* and *VWF* (BEC+LEC), *DCN, LUM* and *PDGFRB* (FRC)*, PROX1* (LEC). **f.** Heatmap showing the top 50 differentially expressed genes at single cell level, marker genes are highlighted. Bars placed above the heatmap indicate the LNSC phenotype: fibroblasts (red), LEC (blue), BEC (green), z-scores expressing fold-change expression compared to mean expression.

Additionally, we confirmed the annotation using differential expression analysis, which revealed a clear signature of the fibroblast, lymphatic and blood endothelial subsets, the top 50 DEGs are represented in the heatmap (Fig.1f; Table 1). In aggregate, this scRNAseq analyses confirmed to presence of LECs, BECs and fibroblasts within our sorted human LNSC sample. We next sought to identify specific cell subsets within each of these three distinct cell types. First, we focused on the fibroblast population in which clustering analysis identified 10 fibroblasts subsets illustrated in the UMAP (Fig. 2a). The number of cells per subset is shown in figure 2b. Additionally, differential expression analysis identified a variety of genes being differentially expressed (DEG) in each subset (Table 2). The top 50 DEGs for each fibroblast subset showing the highest fold change (log2-fold change >0.5, adj-pvalue < 0.05, proportion of cluster expressing > 0.25) are visualized in a heatmap with key genes highlighted for each subset (Fig. 2c). For reference, we assigned a name to each subset based on a selection of key genes highly differentially expressed in each subset (Fig. 2d). Subsequently, the DEGs have been used as input for a GO enrichment analysis reflecting functional differences between each fibroblast cluster subset, indicating a unique role for each subset crucial for the functionality of a lymph node (Fig. 2e). We next compared our identified subsets with subsets previously described in mice and human LNSC studies^5,19–23^ and observed a marked overlap of specific marker-gene lists for the identified fibroblast subsets (Figure 2f). Accordingly, fibroblast subsets identified previously in mice and human studies confirm our findings although nomenclature is sometimes different probably due to differences in sequencing depth. FDC and MRC subsets were not identified in our study, probably due to our sorting strategy. SEPT4+ SC and GLDN+ SC subsets were only identified in our scRNAseq dataset. Next, we elucidated the heterogeneity of the LEC and BEC populations. We applied the same principle for the annotation of the LEC and BEC subsets, as described above for the fibroblast subsets. The analysis of the LECs uncovered four major subsets: ACKR4+ LECs, ACKR1+ LECs, ANGPT2+ LECs and CD24+ LECs (Figure S2a). Unsupervised clustering analysis of the BECs discovered four subsets, namely CA4+ BECs, CDKN1A+ BECs, ACKR1+ BECs, GJA4 BECs (Figure S3a). We provide a detailed description of the LECs and BECs subsets in the "Supplementary Note”. Overall, the identification of these LN endothelial subsets is in line with earlier reports^7,20,24,29,30^.

**Figure 2.**
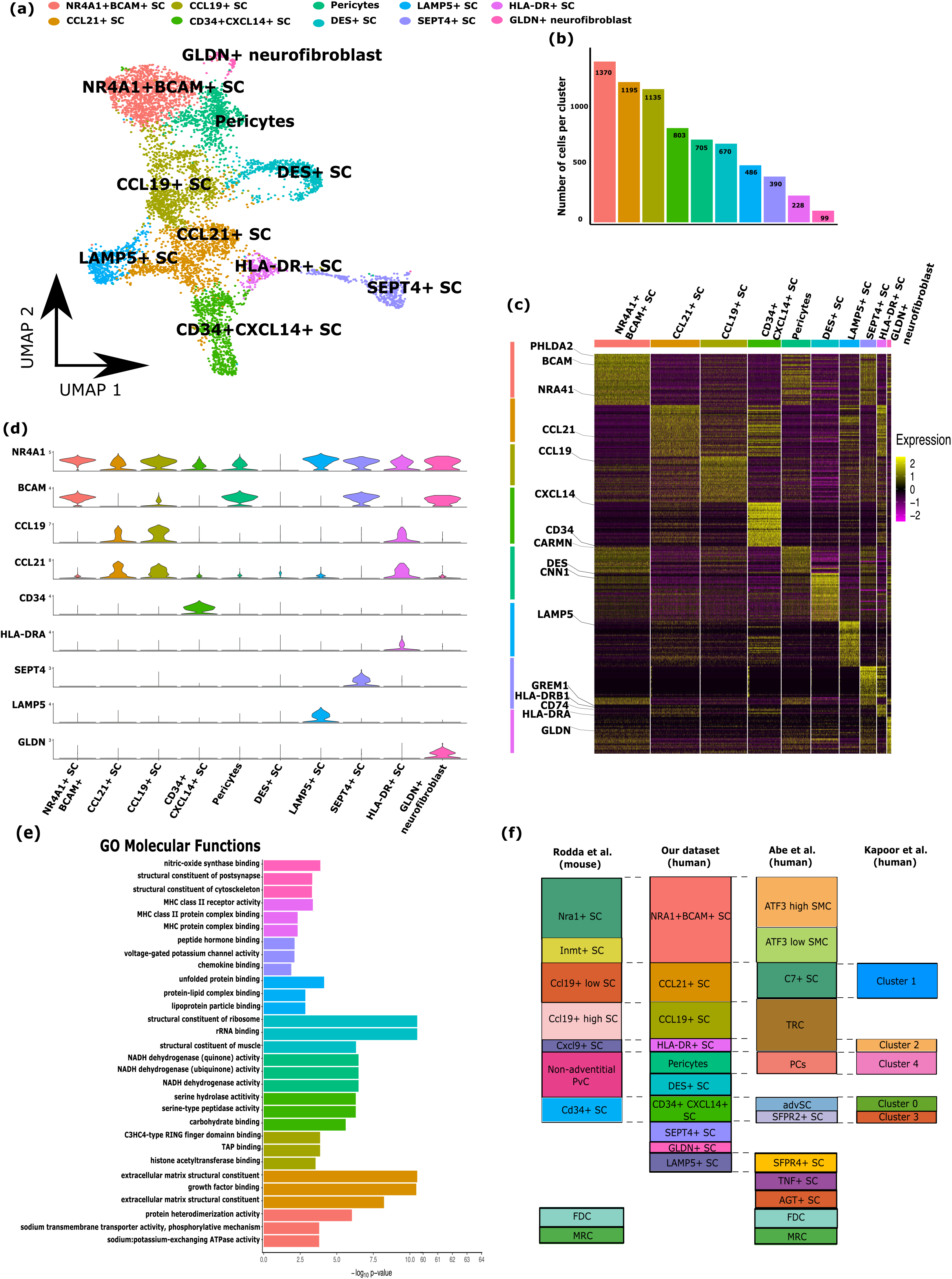
Identification of 10 lymph node fibroblast subsets by scRNAseq. **a.** Uniform manifold approximation and projection (UMAP) analyses revealing 10 fibroblast clusters of LNSCs. **b.** Bar plot indicating absolute count of cells for each cluster. Colors correspond to the cluster to which each cell is assigned. **c.** Heatmap showing per cell (column) the expression patterns of the top 50 DEGs (rows) between the 10 clusters. Marker genes for each cluster are highlighted at the left, at the right, color legend reflecting Z-score expression levels. On top, colored bars to annotate each fibroblast cluster. **d.** Violin plots showing Z-score expression levels of marker genes for each fibroblast cluster. **e.** Gene ontology (GO) enrichment analysis Jusing the DEGs, showing the top 3 GO molecular functions for each fibroblast cluster. **f.** A comparison between the LN fibroblast clusters identified in this study with those previously published by Rodda et al.^19^, Abe et al.^85^, and Kapoor et al^5^. To enable comparison, the bar heights in the previous studies have been adjusted based on the cell numbers assigned to each subcluster in our study.

In summary, this explorative study on a relatively large number of sorted human LNSCs unveiled the presence of 10 subsets of fibroblasts in human LN tissue, of which some have been earlier described e.g. CCL19+ SC, CCL21+ SC, CD34+ SC, DES+ SC, NR4A1+ BCAM+ SC and pericytes, whereas others are newly discovered e.g. SEPT4+ SC, GLDN+ SC, LAMP5+ SC and HLA-DR+ SC. In the following paragraph we further dive into the functionality and spatial location of each fibroblast subset.

### Functional mapping of lymph node fibroblast subsets

We next explored the specific regions within the lymph node potentially occupied by the identified subtypes of fibroblasts described above. We applied a recently developed approach which allows to transfer cell type annotations from scRNA-seq data to a spatial transcriptomic dataset and gives a prediction score for the presence of cells of interest^31^. Hence, we mapped our identified LNSC clusters on a publicly available spatial human lymph node dataset^32^. To first define on a molecular level the LN regions on spot level, we performed cluster analysis of the spatial data and visualised this in a UMAP (Fig. S4a). Secondly, we plotted the cluster identity of each spot of the spatial data, and confirmed that the identified clusters are spatially restricted to specific regions (Fig. 3a). Thirdly, we manually annotated each spatial spot based on the well-established function of genes differentially expressed between specific areas of the lymph node (Fig. 3b, S4b Table 3). Accordingly, we identified canonical regions of the lymph node such as T-cell areas highly expressing *TRBC1*, *TRAC*, follicles expressing *FDCSP*, *CR2,* germinal centres highly expressing *BCL6*, and *MYBL1*, B-T cell interface expressing *THY1,* medulla highly expressing *IGHG1*, *IGHG2*, lymphatic vessels positive for *LYVE1*, *PROX1*, and blood vessels marked by *VWF*, *PECAM1* expression (Fig 3b, S4b, Table 3). Subsequently, we integrated our scRNA-seq dataset with the spatial transcriptomic dataset to obtain for each of the discovered LNSC subsets an *in-silico* prediction score for localisation in each spot of the lymph node tissue spatial transcriptomic dataset (Fig. 3c-n). To provide an objective view of the predicted locations of the identified LNSC subsets per specific region of the lymph node, we calculated for each subset the mean prediction value for every region of the lymph node and visualized this in a heatmap (Fig. 3p). This heatmap shows, for example, that the medulla region which contains many medullary cords are enriched with pericytes and DES+ SC known to surround vessel-like structures. In contrast, the newly identified GLDN+ SC was specifically predicted to locate within germinal center regions of the LN. In aggregate, this spatial analysis, predicting that each identified LN fibroblast subset occupies a different niche within the lymph node, suggests that each LNSC subset harbours a niche-specific function within the LN.

**Figure 3.**
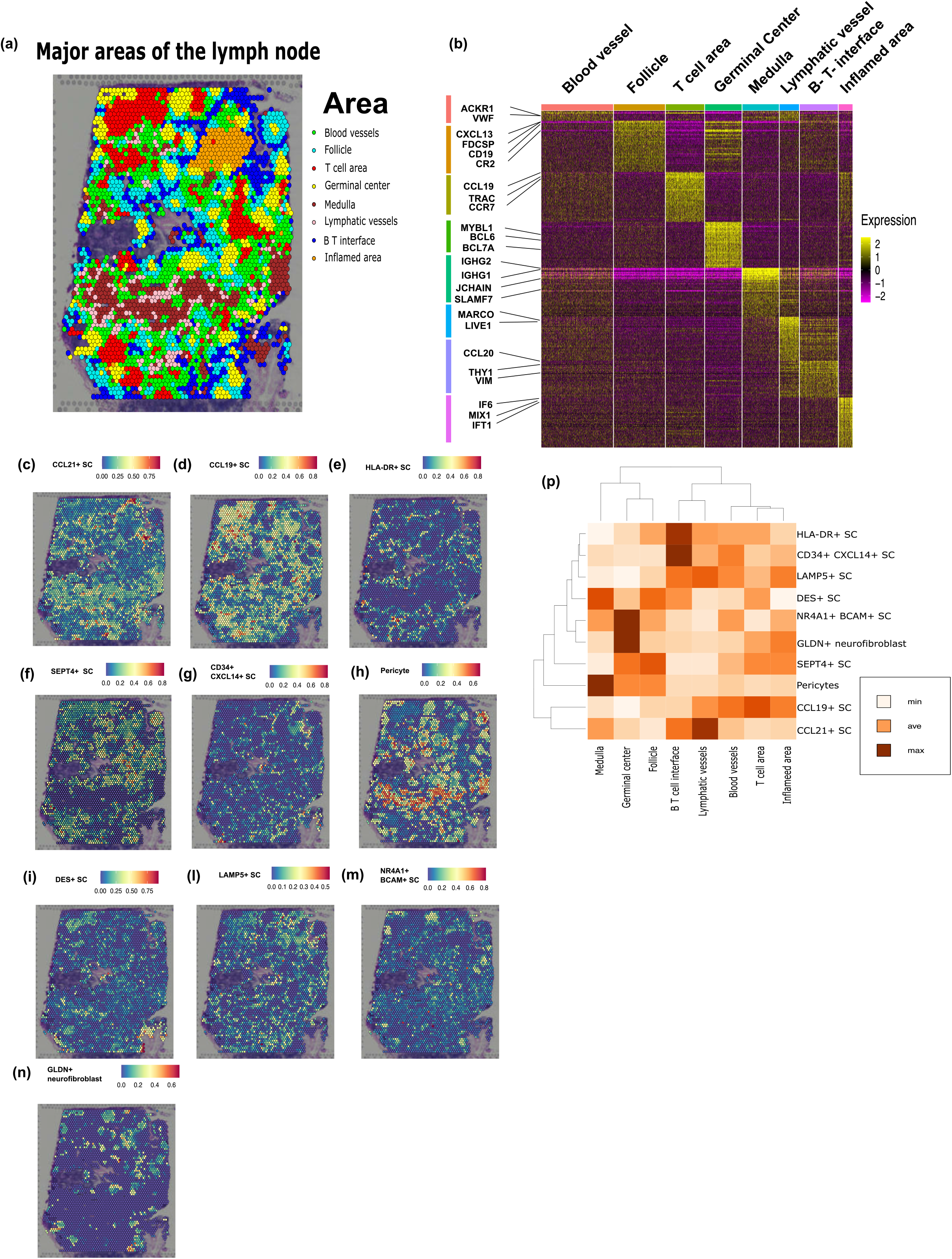
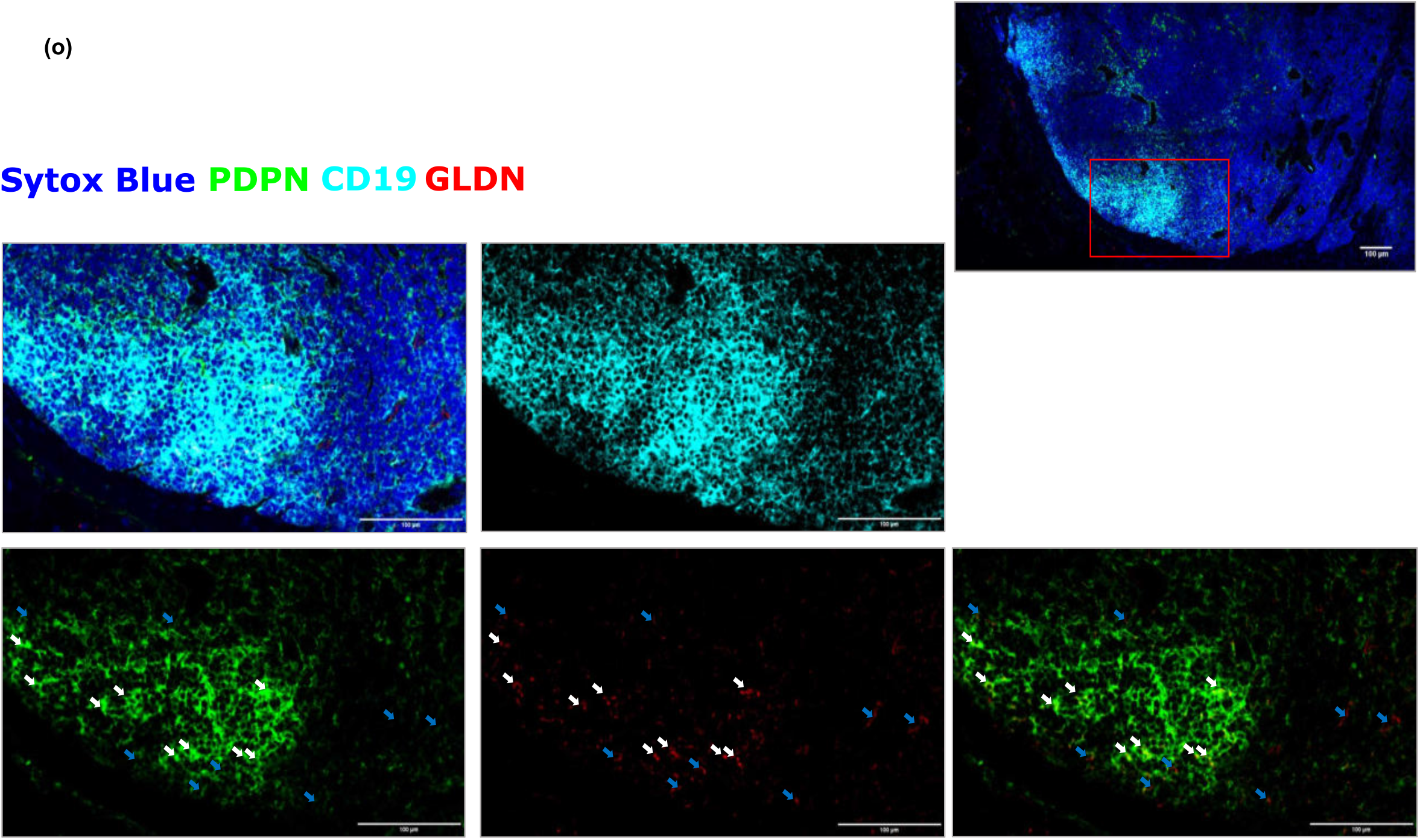
Spatial analyses revealing that different fibroblast subsets occupy distinct regions within the LN. **a.** Spatial gene expression profiles determine major regions within the lymph node: germinal centers, follicles, T-cell and B-cell zones, medulla, blood, and lymphatic vessels (more details in Figure S4). Each region is associated with a different color (see legend). **b.** Heatmap showing differentially expressed genes between different regions of the lymph node which were used to annotate each region (see color bars). At the right, z-scores reflecting expression levels. **c-n.** Visualization of the *in-silico* prediction score per spot for the predicted presence for each fibroblast cluster at the indicated location within the lymph node. Prediction scores range from 0 (blue, not present) to 1 (red, present). **o.** Two-dimensional heatmap clustering the mean values of prediction scores per area of the lymph node for each fibroblast subset. This cluster analysis gives an objective interpretation of the *in-silico* predicted location for each identified fibroblast subset. Mean prediction values range from low, medium, high. **p.** Confocal microscopy images of a human LN tissue section acquired at (40X) stained for PDPN (green), CD19 (cyan), GLDN (red) SytoxBlue (blue). Zoomed image indicate a B cell LN follicle in which PDPN+GLDN+ fibroblasts (white arrows) can be found, while PDPN-GLDN+ are present just outside the follicle. Of note: no staining was observed for neuronal marker NEFL. Representative images from one of three independent experiments are shown.

In the next paragraphs we speculate on the role of these subsets based on DEG, their predicted location, gene ontology analysis and possible interactions with other cells co-localized within the same region of the lymph node.

### CCL19+ and CCL21+ T cell area fibroblast reticular cells

Starting from the cortex of the lymph node, CCL19+ SC also called T-zone reticular cells (TRCs)^19^ expressed high level of CC chemokine ligand 19 (*CCL19*), which guides CCR7+ T cells to the paracortex of the lymph node^33^. In our dataset, CCL19+ SCs express high levels of *CCL2* in the vessel-rich areas of the T cell zone where they might attract monocytes, as previously described in FRCs dwelling the T cell zone^34^. Furthermore, it is worth noting that LNSC predominantly rely on fatty acid metabolism^35^. This is evident through the high expression in CCL19+ SCs of CD36, FABP4, and FABP5, proteins that play crucial roles in the fatty acid metabolism^36^, *FABP4* and *FABP5* also are involved in the maintenance of T lymphocyte homeostasis through the modulation of cytokine production in thymic stromal cells^37^. Additionally, the T-cell zone is populated by the CCL21+ SC, this subset highly express *CCL21* and lower *CCL19* compared to CCL19+ SC. *CCL21* is a chemokine involved in the attraction and homing of the CCR7+ T-cells^38^. CCL21+ SC also highly express stromal cell-derived factor 1 (CXCL12), known for attracting CXCR4+ lymphocytes and monocytes to the lymph node. The gene expression profiles of CCL19+ SC and CCL21+ SC support the mapping analysis which indicated that CCL19+ LNSCs are mostly located in the inner cortex of the T cells whereas CCL21+ LNSCs are predicted to locate more in the outer cortex, closely to the B-T cell interface and in the proximity of lymphatic vessels (Figure 3c, d, p).

### Antigen presenting HLA-DR+ SC and SEPT4+ SC

Our data indicates that HLA-DR+ SC uniquely express high levels of *HLA-DRB1*, *HLA-DRA*, and *CD74*, which are components of the HLA class II antigen presentation machinery^39^. Previous human studies have indeed reported the expression of HLA class II molecules on human LNSCs^40–42^. Furthermore, murine MHC-II+ FRC have been found to be important in regulating antigen-specific CD4+ T cell responses^13,43^. *GREM1* was also highly expressed by HLA-DR+ SC. GREM1+ SC have been recently described as FRC providing a niche for CD4+ T cells in the B-T cell interface of the lymph node and involved in interaction with DC^5^. Our prediction analysis indeed positioned HLA-DR+ SCs in the B-T cell interface (Figure 3e). Based on their transcriptome profile and on their predicted location within the lymph node, human HLA-DR+ SC might engage in self-antigen presentation with CD4+ T cells like their murine counterparts^13,18^. SEPT4+ SCs remarkably express several genes of the septin family including *SEPT7*, *SEPT2*, *SEPT4*, *SEPT9*, *SEPT10*, SEPT11^44^. Septins, in collaboration with actin, organize the filamentous network of the cellular cytoskeleton in dividing and not dividing cells^45,46^, and contribute to autophagosome biogenesis^47^. Our spatial analysis predicts that SEPT4+ SCs is located mostly in the follicles (Figure 3f). We observed that HLA-DR+ SC and SEPT4+SC both highly express septin genes (Table 2) but differ in the expression of genes associated with fibroblast reticular cells as *LUM*, *DCN* and *GREM1*. These findings as well as the UMAP plot may suggest that SEPT4+ SC are functionally related to HLA-DR+ SC, however further investigation is needed to prove this.

### Progenitor CD34+CXCL14+ SC

In the adventitia of the vasculature CD34+ SC may act as progenitor cells that give rise to TRC, FDC, MRC and pericytes^48,49^. As previously reported in mouse, CD34+ SC highly express collagens *COL3A1*, *COL6A2* and *COL1A*2^49^. In accordance with these studies, our mapping analysis assigns high scores for CD34+CXCL14+ SC in the proximity of blood vessels, but even higher unexpected scores for predicted location in the B-T- cell interface area (Figure 3g). The transcriptional profile and predicted location of human CD34+ CXCL14+ SC suggest that this subset indeed may contain progenitor cells giving rise to other lymph node fibroblast subsets, but this warrants further investigation.

### Mural stromal cells

The identified pericytes, DES+ SC, and LAMP5+ SC seem mural cells surrounding vessel-like structures within the lymph node. Accordingly, we observed that pericytes and DES+ SCs have high prediction scores in the medulla (Fig.3h, i), which is rich in medullary cords, whereas LAMP5+ SCs are spatially close to lymphatic and blood vessels (Fig. 3 l, p). Pericytes regulate and support the microvasculature through direct contact with the endothelium^50^. In our dataset pericytes express *MCAM*, *CARMN* and *NET1* consistent with recent studies where human pericytes have been analysed^51–53^. In our dataset they also express *RERGL*, *RGS5*, which have been described as markers for both pericytes and smooth muscle cells in human skin and ovarian cortex^54,55^. Previous studies have described a population of DES+ smooth muscle cells closely wrapping human small muscular arteries^56^. In line with this, our identified DES+ SC highly express genes known to be expressed in smooth muscle cells, including *CNN1, MYL6, MYL9, MYLK, TPM1, TMP2*^57–59^. In our dataset, LAMP5+ SC highly express a variety of genes known to modulate blood pressure and vasculogenesis of the blood vasculature, such as *AGT*, *F5*, *SFPR* ^60–63^. Overall, our study confirms that in the lymph node pericytes and DES+ SC might co-operate with endothelial cells as previously described^19,64,65^. Also, it shows that LAMP5+ SC might be closely associated with the endothelial cells lining the lymph node vasculature.

### NR4A1+ BCAM+ SC resembling an activated stromal cell phenotype in germinal centers

The NRA41+ BCAM+ SCs are predicted to reside in germinal centers (Fig. 3m). As previously described in mouse^19^, also in our dataset NR4A1+ SC highly express *NR4A1*, *FOSB*, *FOS*, *JUNB*, *EGR1*, *NFKBIA* and *ZFP36*. In mouse, this subset is considered to consist of a mixture of different activated cells from other LNSC subsets^19^. We found in our dataset that NR4A1+ SC highly express *BCAM* and *MCAM*, which belong to the laminin family of receptors for extracellular matrix proteins. In murine lymph nodes, FRC-derived laminins have been described to be crucial for localization and transmigration of Th1, Th2 and Th17^66^. In addition, in murine germinal centers, laminin-binding integrins expressed by lymphocytes, promote their migration^67^. The role of human NRA41+ BCAM+ SCs in immunomodulation and lymphocyte trafficking within germinal centers should be further explored to comprehend the functionality of this subset.

### GLDN+ neuro-fibroblasts

Next to genes typically expressed by stromal cells, GLDN+ neuro-fibroblast highly express genes related to neural cells such as *GLDN*, *PCDH10*, *KCNMA1*, ALKAL2, *ADGRL3*^68–73^. *GLDN* and *ADGRL3* are involved in the localization of node of Ranvier and neuron guidance^74,75^. *KCNMA1*, encoding the voltage- and calcium-activated potassium channel^76^, and *PCDH10* is a neural receptor^77^. It is known that in certain circumstances fibroblasts can express neural markers, e.g., human dermal fibroblasts expressing *GLDN*^78^ and in the human brain *FBXO32* expressed by epithelial cells during epithelial–mesenchymal transition^79^. Our mapping analysis revealed that GLDN+ neuro-fibroblasts are located within germinal centers (Fig. 3n, p). Importantly we could validate this finding with immunofluorescence microscopy analysis of human LN tissue sections stained for PDPN, GLDN, and neuronal marker NEFL. Microscopy analyses revealed GLDN+NEFL- neuro-fibroblasts exclusively in the follicles (Fig. 3o). We observed a PDPN+GLDN+ population (Fig. 3o white arrows) in the inner most compartment and PDPN-GLDN+ (Fig. 3o blue arrows) cells in the outmost compartment of follicles. It has been reported in mouse spleen and lymph nodes that interaction between nerve fibres and lymphocytes takes place where humoral immune responses are initiated^80,81^. In the germinal centre of the spatial dataset neuronal markers *TUBB*^82^ and *TUBB4*^83^ are highly expressed, suggesting a high density of neurons in this area of the lymph node (Figure S4c). It will be interesting to study the interaction between human GLDN+ neuro-fibroblast and nerve cells in germinal centre.

### LNSC-immune cell crosstalk

As the data indicates undoubtedly, distinct LNSC subtypes can interact with immune cells at different regions throughout the lymph node. We next explored the possible stroma-specific interaction between T-cells and B-cells, utilizing NicheNet. NicheNet is a powerful framework that enables the identification and analysis of ligand-receptor interactions within complex cellular systems^84^. Notably, this interaction analysis was specifically performed for T-cells interacting with CCL19+ SC, CD34+ SC, HLA-DR+ SC, and B-cells interacting with SEPT4+ SC, NR4A1+ SC. Accordingly, we selected interacting cells well reported in literature i.e., CCL19+ SC with T-cells, but also unknown interacting cells like GLDN+ SC with B-cells. We opted for this strategy because validating findings from well-known interactions would strengthen the validity of thus far unreported interactions. NicheNet predicted several ligand–target and ligand-receptor interactions between GLDN+ SC, NR4A1+ SCs, SEPT4+ SC and B-cells (Fig. 4a, selected targets and receptors in bold) similarly, between CCL19+ SC, CD34+ SC, HLA-DR+ SC and T-cells (Fig. 4b, selected targets and receptors in bold). This analyses clearly shows that the distinct LNSC subtypes interacting with B or T cells, each use distinct ligand-receptor pairs, thereby probably resulting in a different immunomodulatory outcome.

**Figure 4.**
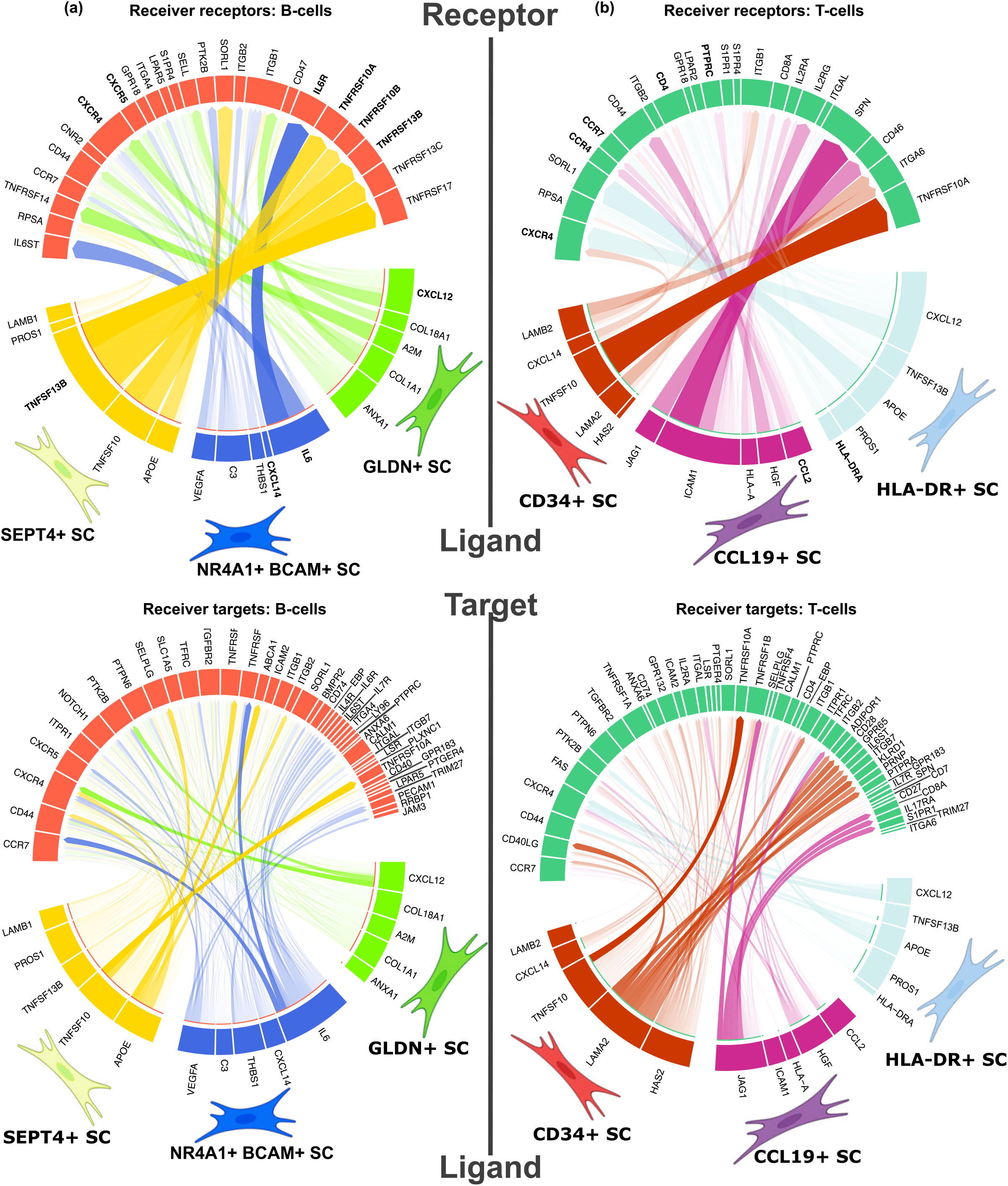
NicheNet to model LNSC-immune cell crosstalk. Circos plots showing the top ligand–receptor (**a, b**) and top ligand-target (**c, d**) pairs identified by NicheNet. Stronger relationships between ligand-receptor or ligand-target pairs are demonstrated by both wider arrow width and lower transparency arrow colors. On the left-side intercellular communication between B cells (red) and SEPT4+ SC (yellow), NR4A1+ SC (blue), or GLDN+ SC (green) is shown. On the right-side crosstalk between T cells (dark green) and CD34+ SC (dark red), CCL19+ SC (purple), or HLA-DR+ SC (cyan) is shown.

## Discussion

In this study we combined our human single cell LN stromal cell data with publicly available spatial data to delineate the heterogeneity of the human lymph node stromal compartment. Our findings reveal a previously unpresented transcriptional, spatial, and functional heterogeneity of human LNSCs. Overall, the predicted spatial organization of LNSCs strongly support the hypothesis that the transcriptional differences across LNSCs subset reflect their distinct functionalities during homeostatic and pathological conditions. Moreover, we show that integrating scRNAseq data with spatial transcriptomics provides a molecular blueprint of the complex heterogeneous stromal architecture within a lymph node. Although the current study is quite unique as it concerns non-disease human LN tissue data, there are some limitations to discuss. The relative low quantity of cells and tissues analysed, may not be sufficient to identify all stromal cell subpopulations. Moreover, further functional experiments are required to validate our findings for each subset.

Even though we sequenced LN stromal cells from only one donor, we could validate our approach by confirming the location of the newly discovered GLDN+SCs by microscopy analysis of human lymph node tissue sections derived from several human donors. Moreover, integrating single-cell RNA sequencing (scRNA-seq) data with a publicly available spatial transcriptomics^32^ corroborated findings that were initially identified through scRNA-seq alone. This underlines the significance of combining spatial and scRNA-seq methods together. Overall, this study provides a data dense blueprint of the human lymph node stromal cell compartment during homeostatic conditions. Our findings indicate that the identified LNSC subsets, characterized by different gene signatures and mapping to specific locations within a healthy human lymph node, each harbour a specialized function which warrants further exploration.

## Acknowledgments

This work was financially supported by grants from NWO-ZonMw (TOP 01217014 to R.E.M. and L.G.M.v.B and VIDI no 91718371 to L.G.M.v.B). We acknowledge the Amsterdam Medical Center Flowcytometry core facility for assistance with the generation of Flow Cytometry data. This work was carried out on the Dutch national e-infrastructure with the support of SURF Cooperative.

## Author contributions

Conceptualization, R.M., L.v.B.; methodology, R.M., L.v.B., P.M., C.G.; investigation, C.G., J.R., R.M., L.v.B., with assistance from C.G.G., J.F.S., A.J., E.R.; formal analysis, C.G. with assistance from P.M., A.J.; resources, E.R.; writing – original draft, C.G., R.M., L.v.B.; writing – review & editing, C.G., R.M., L.v.B., P.M.; supervision & funding acquisition, R.M., L.v.B.

---

**Figure S1.**
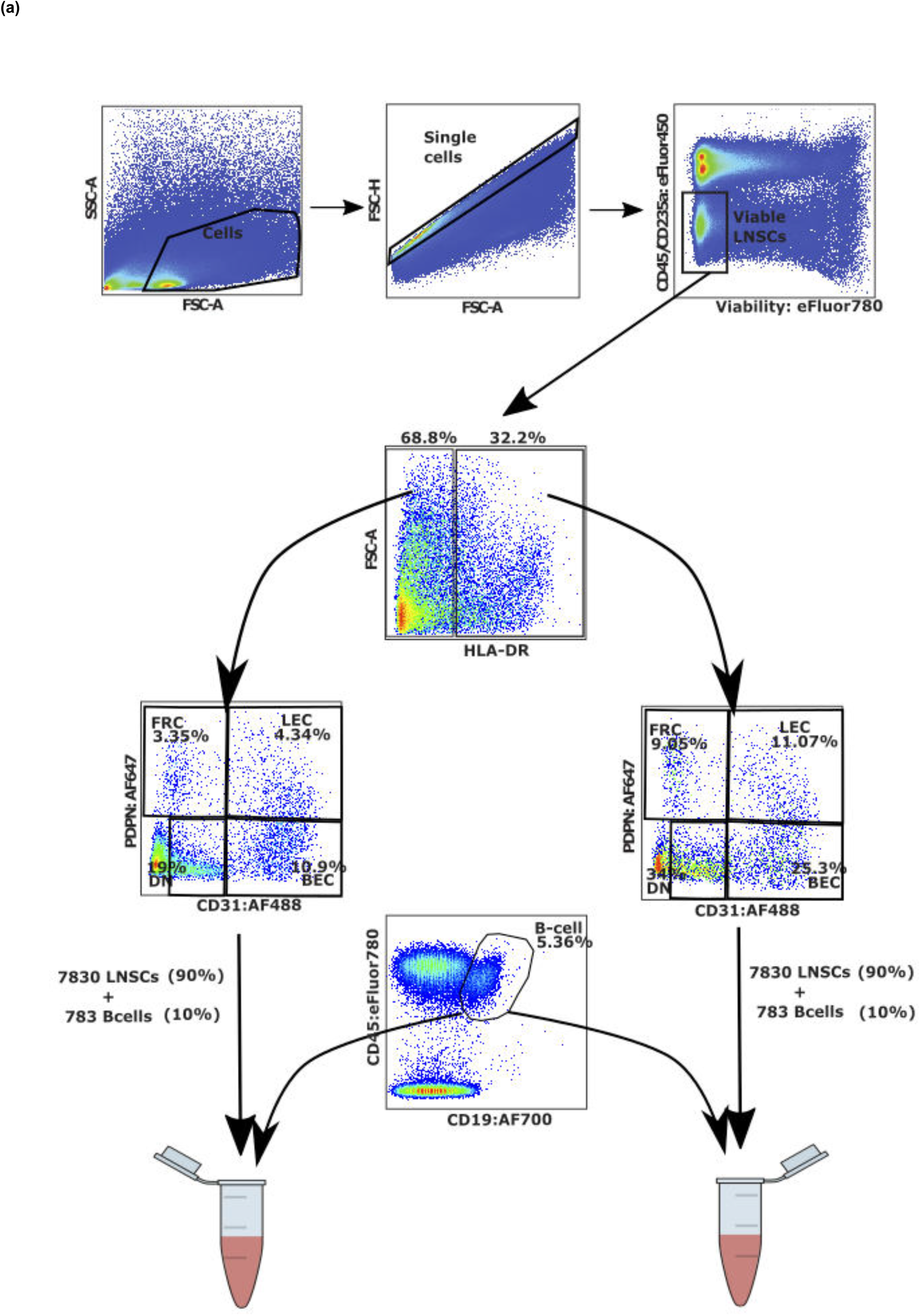
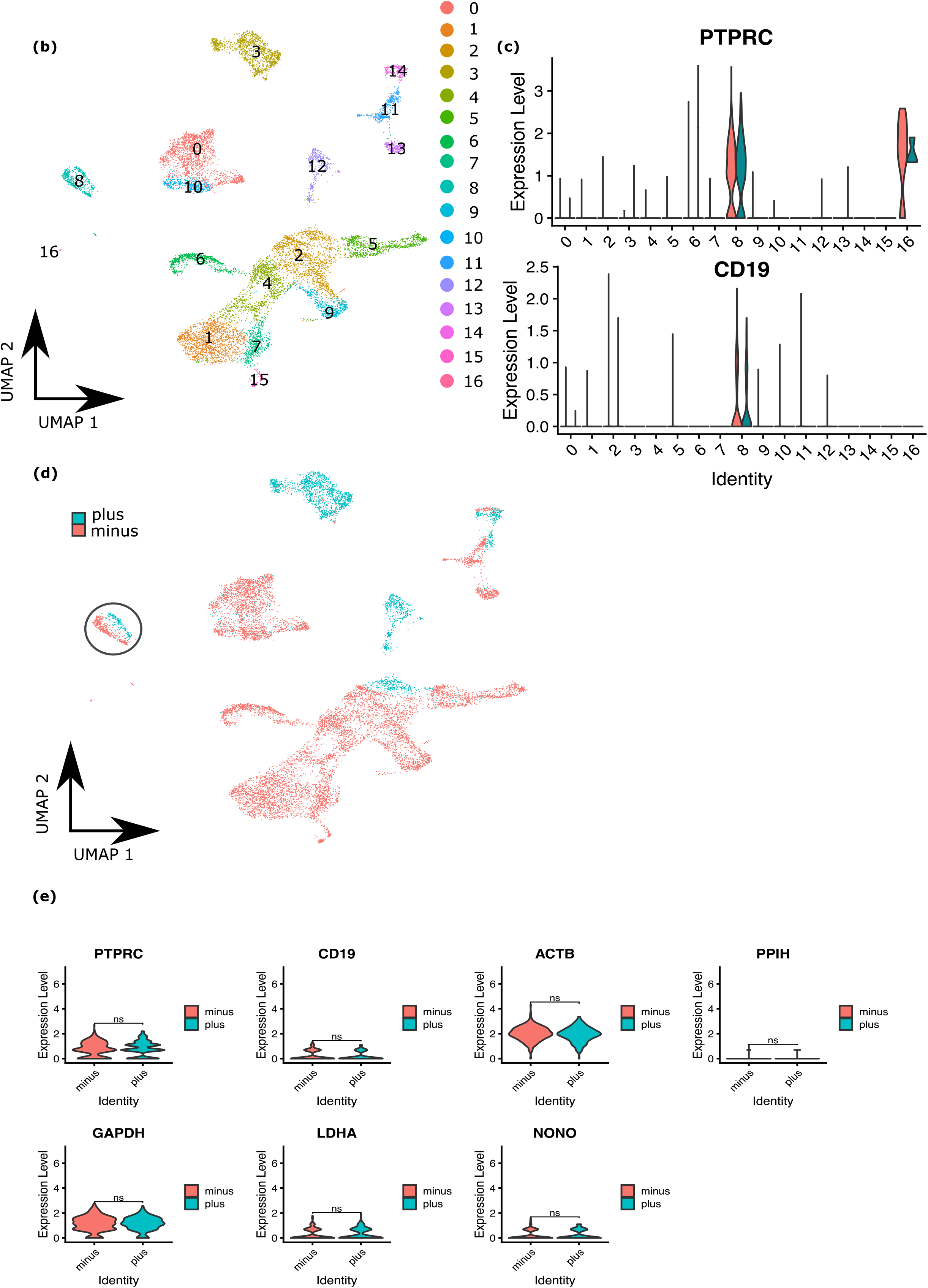
Sorting strategy and 10X Genomics quality control. **a.** Fluorescence-activated cell sorting (FACS) gating strategy used to enrich for CD45-CD235a- viable, single lymph node stromal cells, where we ensured a balance between the major HLA-DR+/- LNSC subsets (DC, FRC, LEC and BEC). In addition, to control for potential 10X Genomics batch effects between the 2 wells, we spiked in B cells from the same sort experiment. **b.** UMAP clustering displaying the clusters present in all sorted cells. **c.** Violin plot showing the Z-score gene expression levels of known B-cell markers *CD19* and *PTPRC* (*CD45*) to determine the clusters containing spiked B cells. **d.** UMAP clustering visualizing the two 10X Genomics wells (plus means sorted HLA-DR+ LNSCs and minus means sorted HLA-DR-LNSCs) and marking the spiked B-cell cluster with a circle. **e.** Violin plots comparing the Z-score gene expression level of several housekeeping genes (*ACTB, PPIH, GAPDH, LDHA, NONO*), and B-cell specific markers (*PTPRC, CD19*) between the two 10X Genomics wells. Statistical significance was tested with Wilcoxon signed-rank test, n.s. (not significant, P-value>0.05).

**Figure S2.**
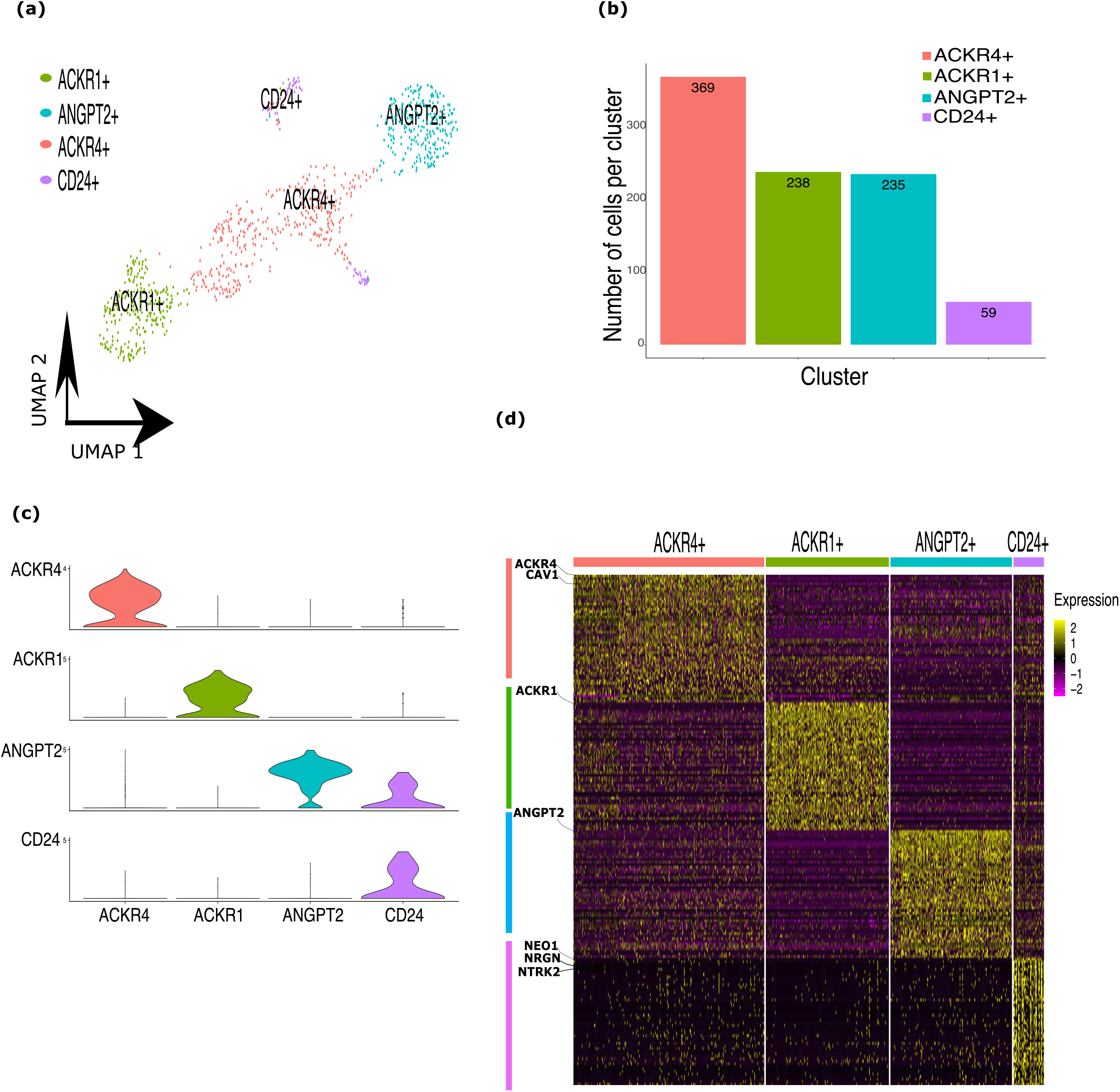
Identification of 4 clusters of LECs in the lymph node. **a.** UMAP plot showing LEC clusters. **b.** Bar plot indicating absolute count of cells for each cluster. Colors correspond to the cluster to which each cell is assigned. **c.** Violin plots representing the Z-score expression levels of top marker genes for each LEC cluster. **d.** Heatmap showing the expression levels of top-50 marker genes for each LEC cluster. Key genes are indicated on the left.

**Figure S3.**
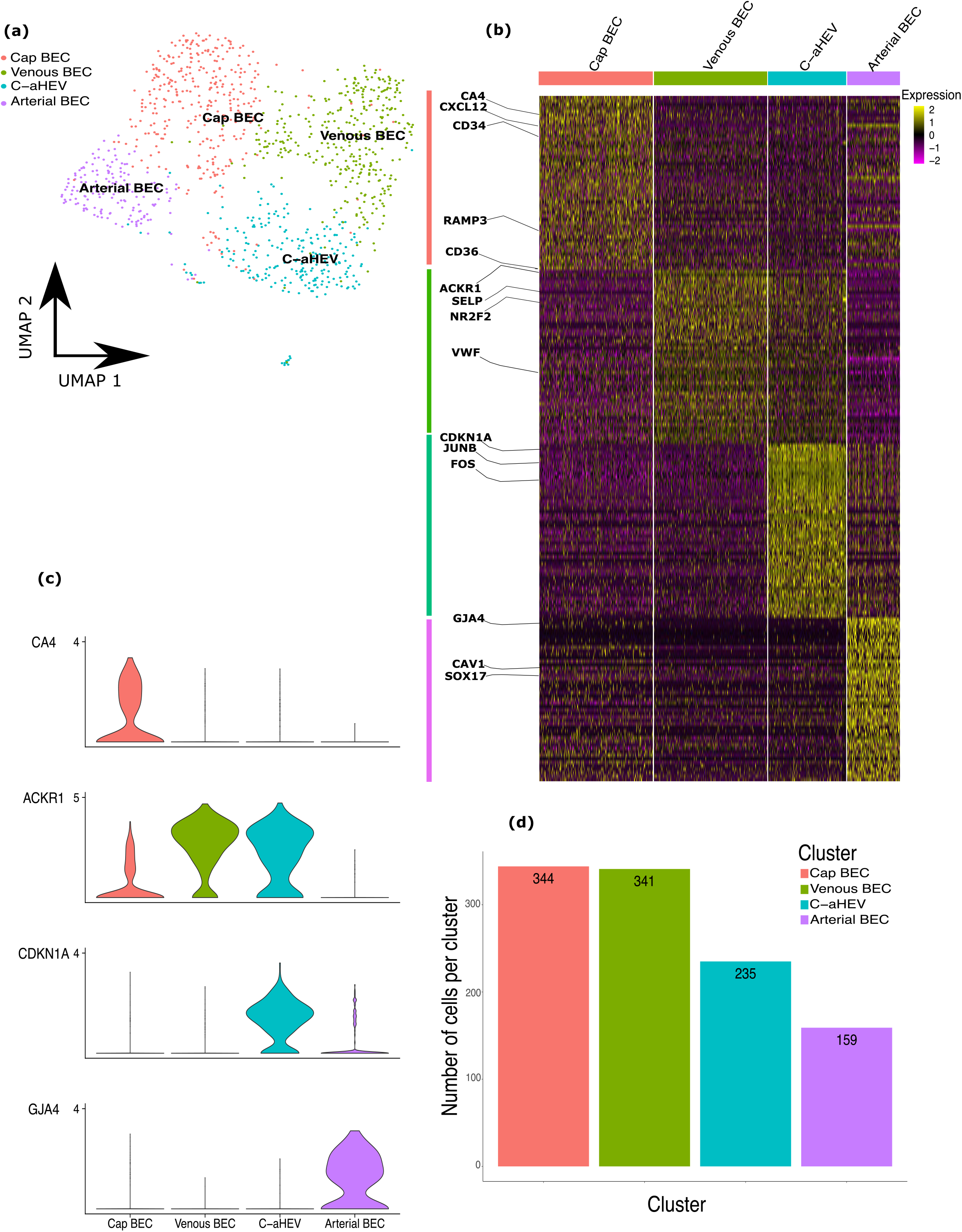
Identification of 4 clusters of BECs in the lymph node. **a.** UMAP plot showing BEC clusters. **b.** Bar plot indicating absolute count of cells for each cluster. Colors correspond to the cluster to which each cell is assigned. **c.** Violin plots representing the Z-score expression levels of top marker genes for each BEC cluster. **d.** Heatmap showing the expression of top-50 marker genes for each BEC subcluster. Key genes are indicated on the left.

**Figure S4.**
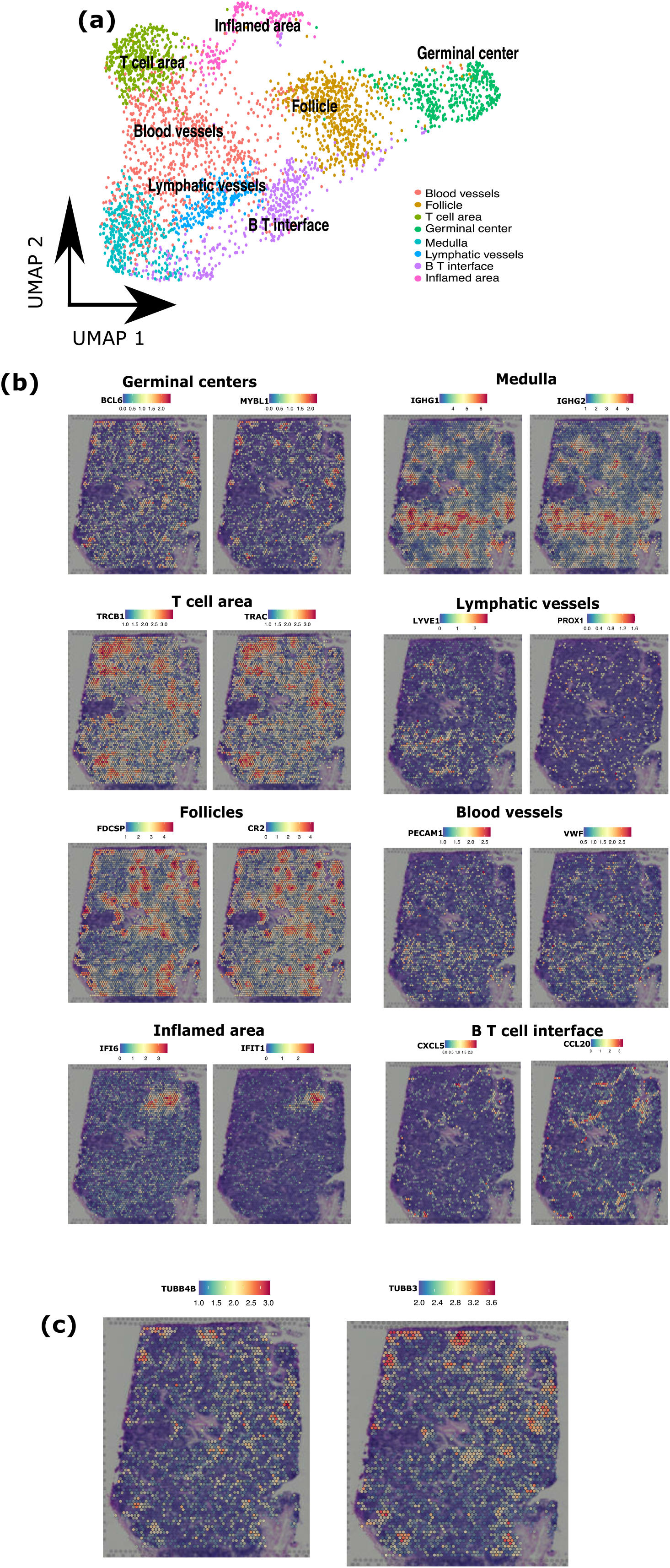
Annotation of main cellular regions on a human lymph node tissue spatial transcriptomic dataset. **a.** UMAP plot showing different lymph node areas (clusters) identified by unsupervised clustering. Clusters were manually annotated based on DEG (see figure 3b). **b.** Spatial plot visualizing expression levels of marker genes (e.g., *BCL6, TRBC1*) known to be highly expressed in specific areas of a lymph node to confirm lymph node area annotations (e.g., Germinal centers, Medulla) or **c.** to visualize marker genes known to be expressed by neurons (*TUBB4B, TUBB*). Z-scores indicate expression level, ranging from 0 (blue) to max (red).

## STAR★Methods

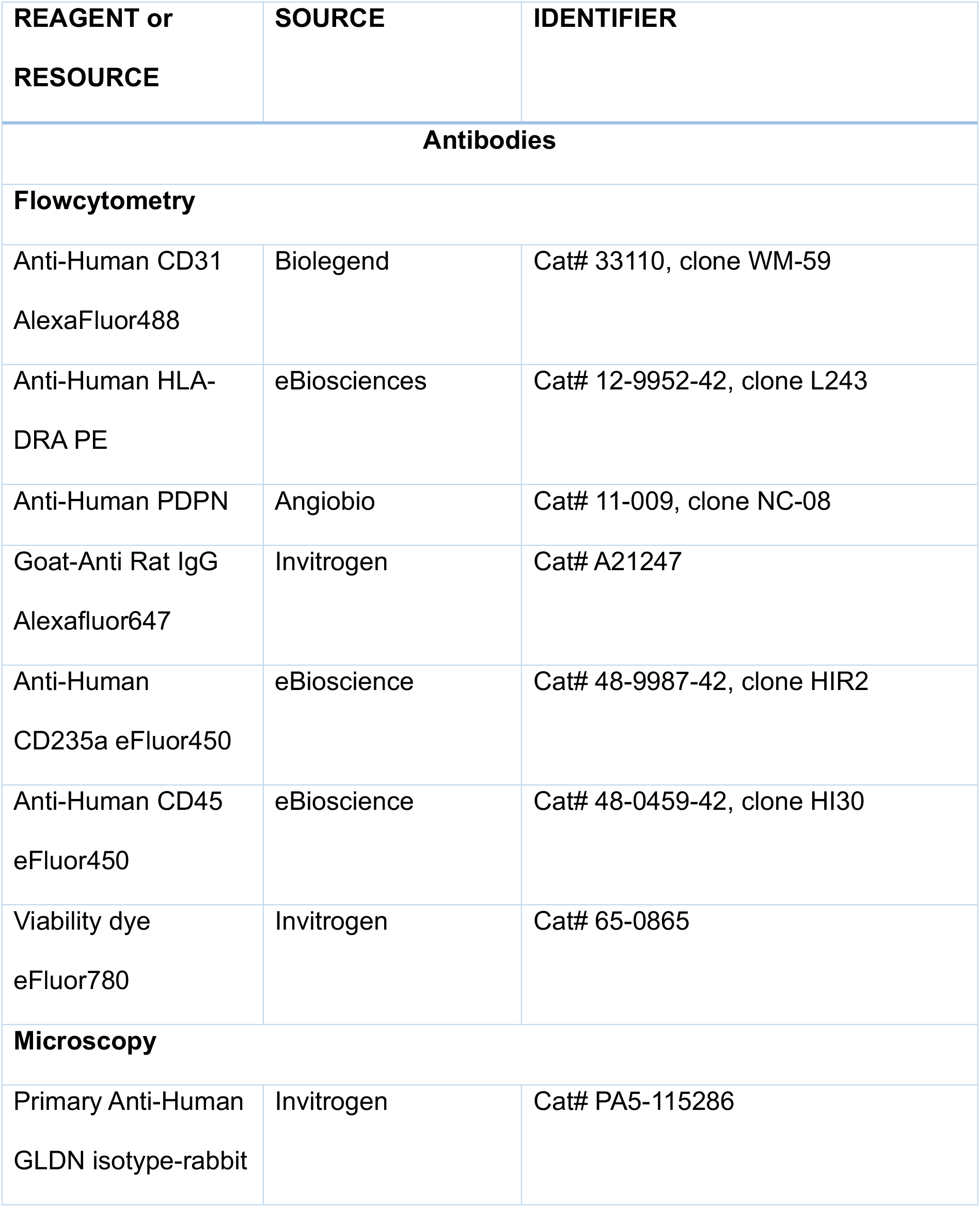
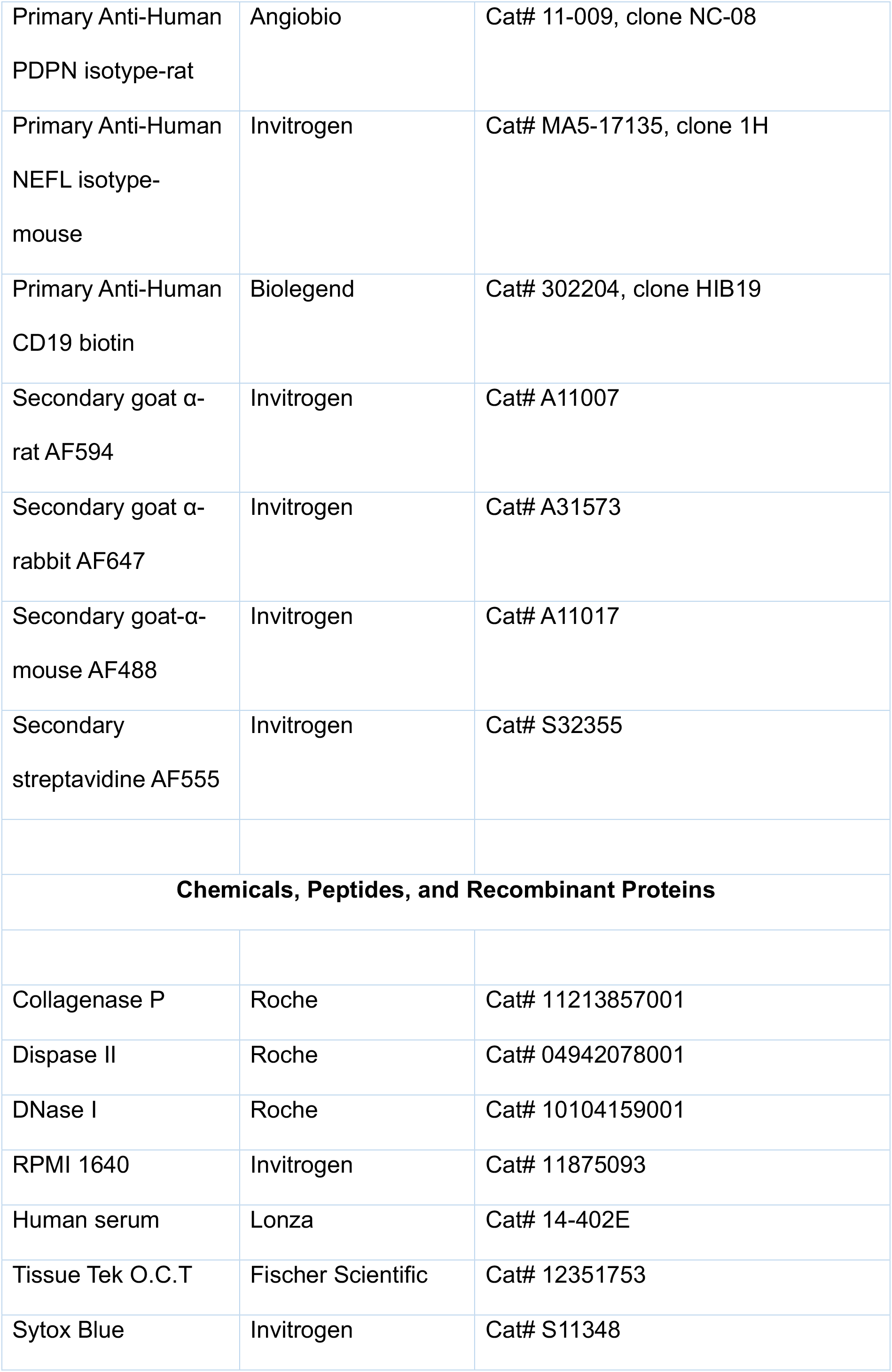
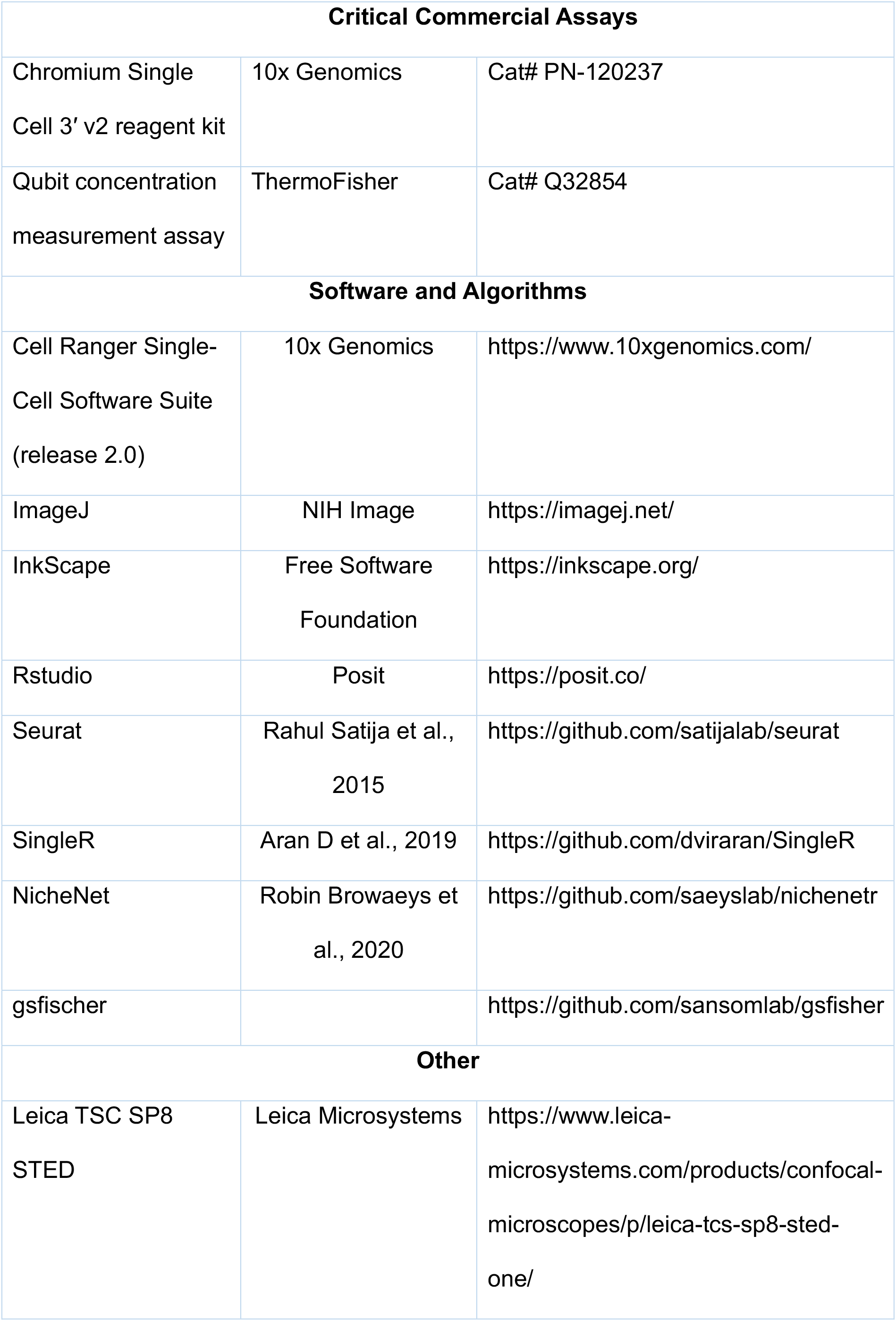
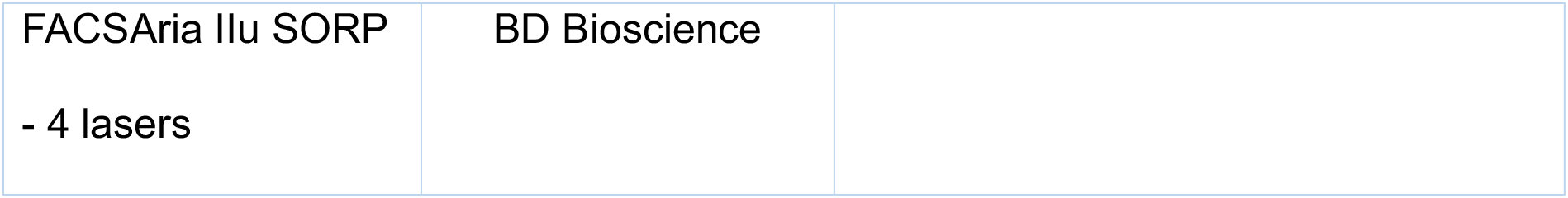

### Stromal Cell Preparation and Flow Cytometry Sorting

LNs were collected from kidney transplantation recipients during living donor kidney transplantation as described before^86^. The tissue did not show any signs of inflammation. As previously described^87^, in short biopsy was digested using the enzymatic mixture of 0.2 mg/ml collagenase P (Roche), 0.8 mg/ml Dispase II (Roche) and 0.1 mg/ml DNase I (Roche) in RPMI medium without serum (Gibco). After, cells were twice washed in PBA buffer (PBS containing 0.01% NaN3 and 0.5% bovine serum albumin (BSA). Cell suspensions was filtered through a 100-μm nylon cell strainer (BD Falcon). Single-cell suspension was stained for sorting flow cytometry using antibodies against CD31 (AlexaFluor488, Biolegend, Cat# 33110, clone WM- 59), HLA-DR (PE, eBiosciences, Cat# 12-9952-42, clone L243), PDPN (Angiobio, Cat# 11-009, clone NC-08), Goat-Anti Rat IgG (Alexafluor647, Invitrogen, Cat# A21247), CD235a (eFluor450, eBioscience, Cat# 48-9987-42, clone HIR2), CD45 (eFluor450, eBioscience, Cat# 48-0459-42, clone HI30), Viability dye (eFluor780, Invitrogen, Cat# 65-0865). Suspension was sorted at the FACSAria IIu SORP - 4 lasers (BD Bioscience). After sorting freshly sorted cells were immediately counted and processed according to 10X Genomics guidelines (CG000126 Guidelines for Optimal Sample Prep Flow Chart RevA). The medical ethics committee of the Academic Medical Center, Amsterdam, approved this study and all subjects gave written informed consent in accordance with the Declaration of Helsinki.

### Library preparation for single-cell mRNA-sequencing

The 2 sorted samples (HLA-DR+ and HLA-DR- LNSCs) were individually sequenced at the core facility after which data was merged. Single-cell RNA-sequencing libraries were prepared according to the manufacturer’s instructions (CG000204 Rev D) using Chromium Next GEM Single Cell 3ʹ GEM, Library & Gel Bead Kit v3.1 (10X Genomics, PN-1000121) and Chromium Next GEM Chip G Single Cell Kit (10X Genomics, PN-1000120). Shortly, cells were combined with reverse transcriptase (RT) Master Mix and partitioned into nanoliter-scale Gel Bead-In-EMulsions (GEMs) using 10X GemCode Technology where the poly-A transcripts are being captured and barcoded with an Illumina TruSeq Read 1 sequence, a 16 bp 10X barcode (unique for each individual cell), a 12 bp Unique Molecular Identifier (UMI; unique for each transcript). GEM generation and Barcoding including fragmentation, end repair and A-tailing, double sided size selection, adapter ligation, clean-up, index PCR, double sided size selection, and QC were performed according to the protocol provided by 10x Genomics (https://support.10xgenomics.com/single-cell-gene-expression/library-prep/doc/user-guide-chromium-single-cell-3-reagent-kits-user-guide-v31-chemistry). Library quality control was performed with the Bioanalyzer HS DNA chip, like the quality check of the cDNA earlier in the protocol. Quantification was performed using a Qubit concentration measurement assay (Quant-it DNA HS Assay kit, ThermoFisher, Q32854). Sequencing was performed with Illumina HiSeq4000 using 2 × 150 bp chemistry.

### Processing raw data from scRNA-seq of 10x Genomics

Demultiplexing of raw base call (BCL) files and conversion to FASTQ files was performed with the Cell Ranger pipeline (10x Genomics, version 3.0.2) using ‘cellranger mkfastq’. Alignment to the human reference genome (10x Genomics; GRCh38 version 3.1.0), read filtering, barcode correction, UMI counting and filtering of empty barcodes was performed using ‘cellranger count’ to generate feature counts for every single cell. The 2 samples were combined using the function ‘merge’ (Seurat), resulting in a dataset containing 13,259 cells. The mean number of UMI counts per cell was 47,582, the median number of expressed genes for each cell was 1,778. To remove cells of low quality, cells expressing less than 300 genes and cells with more than 20% of reads mapping to mitochondrial genes were filtered out.

Genes with at least one feature count in more than three cells were used in the analysis.

### Data analysis of scRNA-seq data

The filtered count data were analyzed using the Seurat package (version 3.2). Global scaling normalization was applied with the ‘NormalizeData’ function via the ‘LogNormalize’ method and a scale factor of 10,000. Next, we apply a linear transformation using ‘ScaleData’ prior to the dimensional reduction. Highly variable genes were identified by ‘FindVariableFeatures’ (selection.method = “vst”) and used for principal component (PC) analysis. The first 50 PCs were used for non-linear dimensionality reduction using UMAP and cell clustering. The resolution of ‘FindClusters’ was set to 0.5 because the resulting clusters were consistent with the mapping of known markers. Reference-based annotation was performed using ‘Blueprint EncodeDataset’ from SingleR^88^ (version 3.12). Non stromal cells were filtered out, when annotated as B-cells, CD4+ T-cells, CD8+ T-cells, HSC, macrophages, melanocytes, mesangial cells, monocytes, NK cells (Table 6). After filtering, the stromal cell dataset was slim down using the function ‘DietSeurat’, followed by linear transformation using ‘ScaleData’, dimensional reduction ‘RunPCA’ (50 dimensions), ‘FindClusters’ (0.5 resolution), resulting in 16 clusters. DEG were identified using the ‘FindAllMarkers’ with the default parameters. We removed cluster 12, and 14 because highly expressing *JCHAI,* which is highly expressed by specific B cells. Next, we performed once more ‘DietSeurat’, ‘ScaleData’, ‘RunPCA’ (50 dimensions), ‘FindClusters’ (0.5 resolution), resulting in 16 clusters. Again, DEG were identified using the ‘FindAllMarkers’ with the default parameters. Based on the expression of know genes we aggregated the 16 clusters in 3 major populations namely fibroblasts, BECs and LECs.

### Fibroblast, BECs, and LECs dataset

Firstly, to obtain the fibroblast dataset, we subset from the stromal cell dataset cluster highly expressing PDGFRB and/or PDGFRA and VWF (CD31) negative. Secondly, for the BECs dataset, we selected from the stromal cell dataset those clusters highly expressing VWF. Thirdly, we selected those clusters highly expressing LYVE and/or PROX1, and VWF. After filtering, the 3 datasets were slim down using the function ‘DietSeurat’, followed by linear transformation using ‘ScaleData’, then the top 2000 variable genes were selected using ‘FindVariableFeatures’ for the dimensional reduction ‘RunPCA’ (50 dimensions). We calculated the UMAP coordinates and clustering (50 dimensions and resolution 0.5) using the functions ‘RunUMAP’ and ‘FindClusters’. DEG were identified using the ‘FindAllMarkers’ with the default parameters.

### Gene set enrichment analysis (GSEA)

GSEA was performed using the R package gsfischer (https://github.com/sansomlab/gsfisher) on the gene expression signatures of each cluster resulting from the Wilcoxon test.

### Spatial transcriptomics dataset analysis

The human lymph node spatial dataset was downloaded from the dataset portal provided by 10x Genomics (https://support.10xgenomics.com/spatial-gene-expression/datasets/1.1.0/V1_Human_Lymph_Node). The dataset contains 4,035 detected spots, 20,239 median UMI counts per spot, 5,999 median genes per spot. Low-quality spots corresponding to spots expressing < 1000 genes or > 20% of mitochondrial-associated genes were filtered out. The filtered count data were analysed by the Seurat package (version 3.2). The dataset was normalized using the ‘SCtransform’ function^89^ The first 50 PCAs were used for UMAP dimensionality reduction and cell clustering. The resolution of ‘FindClusters’ was set to 0.8 because the resulting clusters were consistent with the mapping of known markers in each area of the sequenced section.

### Transfer of labels to the spatial transcriptomics dataset

We applied the ‘FindTransferAnchors’ ‘anchor’-based integration workflow introduced in Seurat v3, that enables the probabilistic transfer of annotations from a reference (our dataset) to a query (spatial transcriptomics) set. Anchors represent pairwise correspondences between individual cells (one in each dataset). These anchors are then used to transfer information from reference to query using ‘TransferData’ (weight.reduction = “pcaproject”). The procedure outputs prediction scores, for each spot, as probabilistic classification for each of the scRNA-seq derived clusters.

### Interaction analysis with NicheNet

NicheNet analysis was adapted from the vignette described at https://github.com/saeyslab/nichenetr. In brief, a Seurat object containing the 10 subsets stromal cell, B-cells and T-cells was generated. As NicheNet requires selected a population as ‘receiver’ and others as ‘sender’. We performed two NicheNet runs, one with T cells (receiver) and CCL19+ SC, CD34+ SC, HLA-DR+ SC (senders), and B-cells (receiver) with GLDN+ SC, NR4A1+ SCs, SEPT4+ SC (senders). To determine the ligands expressed of the sender cell subset, we provided the DEGs of each stromal cell subsets previously computed with Seurat were used, subsequently NichNet extracted genes known to express ligands. Genes expressing receptors were calculate with the function get_expressed_genes() provided by NicheNet. NicheNet analysis was performed based on the vignette to infer receptors and targets and generate a circus plot for the top5 ranked ligand for each sender and their respective receptors/targets. Complete table of ligand receptor, -targets interaction of stroma cell within T-cells Table 7 and with B-cells in Table 8.

### Fluorescent immunohistochemistry staining of frozen lymph node section

Renal lymph nodes were embedded in optimal cutting temperature compound (OCT, Fischer Scientific) and directly frozen in liquid nitrogen. The OCT frozen tissue blocks were cut in 7 µm-thick sections for further analysis. Frozen sections were acetone-fixed for 10 minutes at room temperature (RT). After washing with PBS supplemented with 2% newborn calf serum (nbcs, Lonza, cat# DE14-417EH) (PBS-nbcs), the sections were blocked for 15 minutes at RT with 10% normal human serum (Lonza, cat# 14-402E) in PBS-nbcs to prevent nonspecific binding. Then, sections were incubated with primary antibody diluted in PBS-nbcs for 30 minutes at RT. After three washes, sections were incubated with a secondary antibody diluted in PBS-nbcs for 15 minutes at RT. These staining steps were repeated to stain in sequence GLDN (rabbit, Invitrogen, cat# PA5-115286, clone CRG-L2), PDPN (rat, AngioBio, cat# 11-009, clone NZ-1), NEFL (mouse, Invitrogen, cat# MA5-17135, clone 1H3) and CD19 (biotin, Biolegend, cat# 302204, clone HIB19). Secondary antibodies used were goat-α-rat (AF594, Invitrogen, cat# A11007), donkey- α-rabbit (AF647, Invitrogen, cat# A31573), goat-α-mouse (AF488, Invitrogen, cat# A11017) and streptavidine-AF555 (Invitrogen, cat# S32355). When necessary to block nonspecific binding of the secondary antibody, 2% normal rat serum or 2% normal rabbit serum (produced in house, in collaboration with the Amsterdam Animal Research Center (AARC)) was added to the secondary antibody staining.

Furthermore, before CD19 staining, the sections were blocked with 20% normal mouse serum (produced in house, in collaboration with the AARC) in PBS-nbcs for 15 minutes at RT. Lastly, nuclei were counterstained with Sytox Blue (Invitrogen, cat# S11348) and mounted with Mowiol (Sigma Aldrich, cat# 81381). Images were acquired with the Leica TCS SP8 STED 3x confocal microscope (Leica microsystems, IR GmbH) and analysed with ImageJ^90^.

### Visualization

Log-normalized gene expression data was used for visualizations with violin plots (VlnPlot), UMAP plots (FeaturePlot). Scaled log-normalized gene expression data was used for visualizations with dot plots (DotPlot), heatmaps (DoHeatmap) We plotted proportions with barplot (Dittoplot). Prediction scores were displayed with “SpatialFeaturePlot”.

### Quantification and Statistical Analysis

#### Data and Code Availability

scRNA-seq data will be deposited in the GEO database and available upon request.The analysis was performed with R version 4.0.1 (2020-06-06) running under Windows 10 (x64), main R packages used Seurat (3.2), SingleR (3.12), NicheNet (1.1.0)

