## Supplemental Information for "Identification and mapping of human lymph node stromal cell subsets by combining single-cell RNA sequencing with spatial transcriptomics"

### Heterogeneity of human LECs and BECs

The endothelial cells (ECs) that line the lumens of blood and lymphatic vessels play an integral role in the regional specialization of vascular structure and physiology[1], but also control the access of soluble molecules and subcellular particles (including viruses) to the conduit system that guides them to dendritic cells residing in the LN cortex[2]. The analysis of our dataset confirmed previously described four subsets of LECs: ACKR4+, ACKR1+, ANGPT2+ and CD24+[3–5] (Figure S2a). ACKR4+ LECs represent the largest group of LECs in our dataset (Figure S2b).

Based on highly expressed genes we have annotated each subset, the list of genes is illustrated in a violin plot (Fig. S2c) Differential expression analysis shows a defined gene signature for every subset as shown in the heatmap (Figure S2d, Table 4). ACKR4+ LECs abundantly express *ACKR4* and *CAV1*[2]. Same pattern of expression was described previously in human by Takeda on LECs of the ceiling of subcapsular sinus SCSs and afferent collecting lymphatic vessels[5]. *ACKR1* is up regulated in ACKR1+ LECs and has been reported to be expressed on the so-called pre collecting vessels[6], which connect the lymphatic network in-between the capillary vessels and collecting vessels [7,8]. ACKR1+ LECs expressed *CLEC4G* which is in line with a previous report describing that human[9]. *ANGPT2* is highly expressed on ANGPT2+ LECs, which is a ligand of the endothelial tyrosine kinase receptor (*Tie2*). In mouse, Xiang and colleagues address ANGPT2 as a marker for the subset Ptx3-LEC involved in LN remodelling[10]. ANGPT2+ LECs closely resembles the LEC V subset previously described by Takeda and colleagues[9]. In CD24+ LECs, we found a prominent expression of *CD24* which was recently described as marker for the LECs on the upstream side of valves[11]. Also, CD24+ LECs highly express the neurotrophic receptor *NTRK2*), which promotes lymphoid tissue neovascularization[12]. In the lymph node, the blood vessel network is crucial for immune cell trafficking as well as supplying and clearance of nutrients, and metabolites between blood and lymphoid tissue. In our dataset, unsupervised analysis discovered four subsets of BECs, namely Cap BEC BECs, CDKN1A+ BECs, ACKR1+ BECs, GJA4+ BECs (Fig. S3a), which have been previously described in mouse and human[11,13]. We calculated DEGs in each subset to explore the molecular profile of BECs (Fig.S3b, Table 5). We next annotated the clusters based on marker genes from literature[11,13] (Fig. S3c). As previously described, CA4+ BECs or capillary BECs (cap BECs) represents the microvasculature innervating the lymph node. This cluster embrace capillary endothelial cells previously described in mouse as CapEC, CapEC, CapEC2 and the express their known markers e.g.*CA4, CXCL12, CD34, RAMP3*[13,14]*.* Also highly express the glycoprotein *CD36*, which is known to be expressed on microvasculature[15]. CDKN1A+ BECs or activated capillary HEV (C-aHEV) resemble a subcluster of the HEV. As previously described this subset express stress related heat shock proteins (*HSPA1A, HSP90AB1, HSP90AA1*) and JUNK activation proteins (*JUNB, FOS*)[11] ACKR1+ BECs or venous BECs, and GJA4+ BECs or arterial BECs express markers previously described in these subsets, respectively ACKR1 and GJA4[16,17]. We observed that Cap BEC comprise the highest proportion of BECs in our dataset (Fig. S2d). In summary, our scRNAseq analysis of human LN endothelial cells confirm the existence of previously identified LECs[2,5] and BECs[13,18] subsets thereby strengthening these findings.

**Sorting strategy and B-cell spike-in**

We have noticed that HLA-DR+ stromal cells represent only a small population within the stromal compartment of the lymph node. To ensure that sufficient HLA-DR+ stromal cells were included in the sorted cells used for sequencing, we included HLA-DR in the gating strategy of the sorting. We first gated on the HLA-DR+ (plus) and HLA-DR- (minus) stromal cells, subsequently we sorted DN, FRC, BEC and LEC based on PDPN and CD31 expression, resulting in two sorted cell suspensions. To assess the presence of a possible batch effect between these two samples, we added spike-in B cells in each sample (Figure S1a). See Figure S1a for a detailed overview of the gating strategy. After sorting, cells were manually counted and loaded in 2 different wells of the chromium chip. The 2 samples were individually sequenced and then merged. Refer to the method ("Library preparation for single-cell mRNA-sequencing") for more specifications. Spike-in B cells were identified as cluster 8 of the annotated dataset (see Methods) as highly expressing CD19 and PTPRC (CD45), containing cells from both the "minus" and "plus" sort, and made up almost exclusively of cells annotated as B cells (Figure S1b, S1c). To investigate the effects of the experimental design at the level of individual genes, we compared the expression of several housekeeping genes[101], indicating no significant differential expression between "minus" and "plus" sorted B-cells (Figure S1e). Based on these quality checks, we concluded that the experimental design did not introduce any batch effect in the dataset and samples could be merged and analysed.
